# Reducing phytate, an antinutritional factor, in sorghum dough using phytase-expressing *Saccharomyces cerevisiae*

**DOI:** 10.1101/2024.11.13.623345

**Authors:** Tanaya Kale, Rutuja Kamble, Aditi Pawar, Sahil Dawankar, Shamlan M. S. Reshamwala

**Author notes:** Equal contribution (Tanaya Kale, Rutuja Kamble). Corresponding author Address: Department of Biological Sciences and Biotechnology, Institute of Chemical Technology, Nathalal Parekh Marg, Matunga (East), Mumbai 400019, Maharashtra, India.

## Abstract

Phytate, a major antinutritional factor present in millets and legumes, chelates minerals like calcium, iron and zinc leading to their poor bioavailability. Phytase from *Aspergillus japonicus* was expressed extracellularly in *Saccharomyces cerevisiae* and its ability to reduce phytate in sorghum flour was determined. When the *A. japonicus* phytase-expressing strain was used to ferment sorghum dough, phytate reduction increased by 43% as compared to control strain. This approach of simultaneous proofing and phytate reduction can be used to increase micronutrient availability of non-wheat food products.

## 1. Introduction

A major nutritional limitation of wheat bread, an inexpensive staple food, is its low content of essential amino acids, particularly lysine. Moreover, there is a heightened interest in using non-wheat flour for foodstuff preparation due to an increase in gluten-related disorders (Tovoli et al., 2015). Apart from animal sources, which have a protein content twice that of cereals, legumes and millets have emerged as an economical and environmentally sustainable protein source to potentially improve the nutritional value of breads. The use of legume and millet flours to supplement a staple food like wheat bread has immense potential, particularly in developing countries to increase the dietary intake of high-quality protein (Kouris-Blazos and Belski, 2016; Hassan et al., 2021). Legumes are a rich source of essential amino acids, including phenylalanine, leucine, isoleucine, arginine, tryptophan and lysine; moreover, consumption of legumes is associated with reduced risk of colorectal cancer (Kouris-Blazos and Belski, 2016). Similarly, including millets in the diet has a positive effect on human health (Hassan et al., 2021). Therefore, partial replacement of wheat flour by non-wheat flour is one of the common techniques to formulate functional breads (Chavan and Kadam, 1993). In the Asian and African context, as traditional diets contain a variety of millets, supplementation of breads and other bakery products with millet flour may gain greater acceptability.

On the other hand, the presence of antinutrient factors such as phytate have been reported frequently in millets, which limits their application in food products (Lv et al., 2017; Boncompagni et al., 2018). Phytates can reduce protein and amino acid digestibility by up to 10% (Gilani et al., 2005) and can reduce mineral and trace element bioavailability (Schlemmer et al., 2009). Therefore, phytate content has to be reduced to enable unlocking of benefits of using millet flours without the additional drawbacks.

*Sorghum bicolor*, commonly known as sorghum, is a widely consumed millet in Africa and India. The high phytate content in sorghum is associated with low mineral uptake, as phytates chelate the cations and form insoluble complexes that cannot be digested (Rebellato et al., 2020). In the present study, phytase was expressed extracellularly in *S. cerevisiae*, and the constructed strain was used to reduce phytate content and simultaneously proof sorghum flour.

## 2. Materials and Methods

### 2.1 Strains, media and cultivation conditions

*Escherichia coli* DH5α was used as the cloning host and was routinely cultivated in Luria Bertani medium at 37°C. Ampicillin (100 µg/ml) was added when required. The *E. coli*-yeast shuttle vector pRS426-GPD (Mumberg et al., 1995) was used to clone and express genes in *S. cerevisiae* CEN.PK2-1D. The YIplac204 integrative plasmid (Gietz and Sugino, 1988) was used for genome integration in *S. cerevisiae. S. cerevisiae* was routinely cultivated in Yeast Extract Peptone Dextrose (YPD) medium. *S. cerevisiae* strains transformed with episomal and integrative plasmids were selected on Ura and Trp drop out medium, respectively.

### 2.2 Phytase gene construct

The phytase gene (*phyA*) from *Aspergillus japonicus* BCC18313 (GenBank accession EU786166) was codon optimized for expression in *S. cerevisiae*. The native signal sequence was replaced with the α-mating factor signal sequence followed by a glycine-serine linker. The gene was synthesized with flanking BamHI and EcoRI restriction endonuclease recognition sequences (Twist Bioscience).

### 2.3 Cloning of phytase gene

The phytase gene was amplified using primers Fwd_Aj (5’-GC<u>GGATCC</u>ATGAGATTCCCATCCATTTTCACCG-3’; BamHI site underlined) and Rev_Aj (5’-GC<u>GAATTC</u>TCAAGCGAAACATTCAGCCC-3’; EcoRI site underlined) using high fidelity DNA polymerase and cloned in plasmid pRS426-GPD. T4 DNA ligase was used for ligation, and the ligation mixture was transformed in CaCl_2_-competent *E. coli* DH5α. Transformants were selected using ampicillin and verified using colony PCR. The isolated recombinant plasmid was transformed in *S. cerevisiae* CEN.PK2-1D using the lithium acetate-mediated transformation method. Transformed yeast cells were selected on Ura drop out medium.

### 2.4 Genome integration of phytase gene

For genome integration, the YIplac204 integrative plasmid was used. The GPD promoter – phytase gene – transcription terminator sequence was amplified using primers Fwd_RS (5’-GCTGGAGCTCAGTTTATCATTATC-3’) and Rev_RS (5’-CACTATAGGGCGAATTGGG-3’) using the recombinant pRS426-GPD plasmid containing the *phyA* gene sequence as a template. The amplified fragment was gel purified and blunted using Phusion DNA polymerase and ligated into SmaI-digested YIplac204 vector. The ligation mix was transformed in chemically competent *E. coli* DH5α cells. Transformants were selected on LB plates containing ampicillin and successful cloning was confirmed using colony PCR.

Yeast transformation of YIplac204 vector containing the *phyA* expression cassette was carried out using the lithium acetate method. Transformants were selected on Trp drop out medium.

### 2.5 Phytase assay

Phytase enzyme in yeast culture supernatants was assayed using the following protocol. The reaction mixture contained 10 µl culture supernatant and 100 µl 5 mM sodium phytate in 0.5 M sodium acetate buffer (pH 5.0). The reaction mixture was incubated for 30 min at 50°C. The reaction was stopped by adding 500 µl AAM solution (5 N sulfuric acid, 10 mM ammonium molybdate and 100% acetone in a ratio of 1:1:2). After incubating for 1 min, 50 µl 1 M citric acid was added to the reaction, centrifuged at 5000 rpm for 5 min, and the resulting colour was measured at 355 nm (Heinonen and Lahti, 1981).

### 2.6 Sorghum dough fermentation

*S. cerevisiae* strain AP1, expressing phytase from a genome integrated gene (empty YIplac204-integrated strain was used as control) grown in 300 ml YPD broth was harvested by centrifugation and washed twice with deionized water. The wet cell weight of the cell pellets was determined, and cells were resuspended in 10 ml deionized water. Sorghum flour (bought from the local market) was mixed with 2% (w/w of dough) wet mass of the control and PhyA-expressing yeast cells, respectively, and made into a dough with appropriate quantities of water. The dough was fermented for 8 h and the samples were taken before and after the 8 h interval. The samples were dried at 80°C and pulverized for analysing the phytate content (Arjmand et al., 2023).

### 2.7 Estimation of phytate in fermented dough

To measure phytic acid, the samples (0.5 ± 0.05 g) were added to 10 ml of 2.4% (0.64 N) hydrochloric acid and vortexed vigorously for 10 s. The samples were left overnight on a platform shaker at 300 rpm at room temperature. Then, they were centrifuged at 3000 rpm at 10°C for 20 min. The supernatant was filtered through Whatman filter paper into tubes with previously weighed NaCl (1.0 ± 0.05 g). The mixture was vortexed for 30 s for the salt to dissolve and then shaken at 3000 rpm for 20 min. The samples were allowed to settle at −20°C for 20 min, and then centrifuged at 3000 rpm at 10°C for 20 min. 1 ml of the supernatant was diluted to 25 ml with deionized distilled water; 750 µl of this diluted sample was mixed with 250 µl of the Wade reagent (0.03% FeCl3·6H_2_O + 0.3% sulfosalicylic acid) and thoroughly mixed by vortexing. It was then centrifuged at 3000 rpm at 10°C for 10 min, and the absorbance was recorded at 500 nm (Darambazar, 2018).

A standard curve was prepared by reacting a series of standard solutions containing 0, 20, 40, 60, 80 and 100 mg/l of sodium phytate with the Wade reagent. The phytic acid content of the samples was expressed as milligrams of phytic acid per 100 g of sample.

## 3 Results

### 3.1 Expression of phytase in S. cerevisiae

PhyA phytase from *A. japonicus* BCC18313 exhibits high affinity for phytase and exhibits broad pH stability from 2.0 to 8.0 (Promdonkoy et al., 2009). Promdonkoy et al. (2009) expressed this enzyme in *Pichia pastoris* and demonstrated that it can efficiently hydrolyze phytate in corn-based animal feed. We therefore chose to express this phytase in *S. cerevisiae* and evaluate the potential of an *A. japonicus* PhyA-expressing strain to decrease phytate content in sorghum dough.

The *A. japonicus* phytase gene sequence was codon-optimized for expression in *S. cerevisiae*. The native signal sequence was replaced with the α-mating factor signal sequence to allow the enzyme to be secreted in the medium. The gene was cloned in plasmid pRS426-GPD, an episomal vector, under control of the constitutive *GPD* promoter. Expression of phytase was verified by carrying out phytase enzyme assay using cell-free supernatant. Phytase activity was found to be 1080.67±106.65 IU/l/OD_600_, while phytase activity in the empty plasmid containing control strain was 388.88±100.77 IU/l/OD_600_.

To construct a plasmid-free phytase expressing strain, the phytase gene expression cassette, comprising of the CDS flanked by the *GPD* promoter and *CYC1* transcription terminator, were cloned in plasmid YIplac204, an integrative plasmid that integrates in the *trp1* locus and makes the integrant a tryptophan prototroph. Phytase activity of the supernatant of the integrant, called strain AP1, was found to be 1602.78±161.64 IU/l/OD_600_, while the control strain had an activity of 360.89±31.99 IU/l/OD_600_. The increased phytase activity in the integrant as compared with the strain expressing the phytase gene from an episomal plasmid may be due to the medium used: the episomal plasmid constructed strain was cultivated in minimal dropout medium, while the integrant was cultivated in complex medium.

### 3.2 Phytate reduction in sorghum dough

*S. cerevisiae* AP1, expressing phytase from a genome-integrated copy of the phytase gene, was used to ferment sorghum dough. A strain with the empty YIplac204 plasmid integrated in the genome served as control. The dough was allowed to ferment for eight hours at 30°C, and phytate content before and after proofing was determined.

We observed that while phytate content was reduced by 29.35±6.42 mg by the control strain, phytate content was reduced by 42.03±7.92 mg by *S. cerevisiae* AP1, representing a 43% increase in phytate reduction (P=0.017; Fig. 1).

**Fig. 1.**
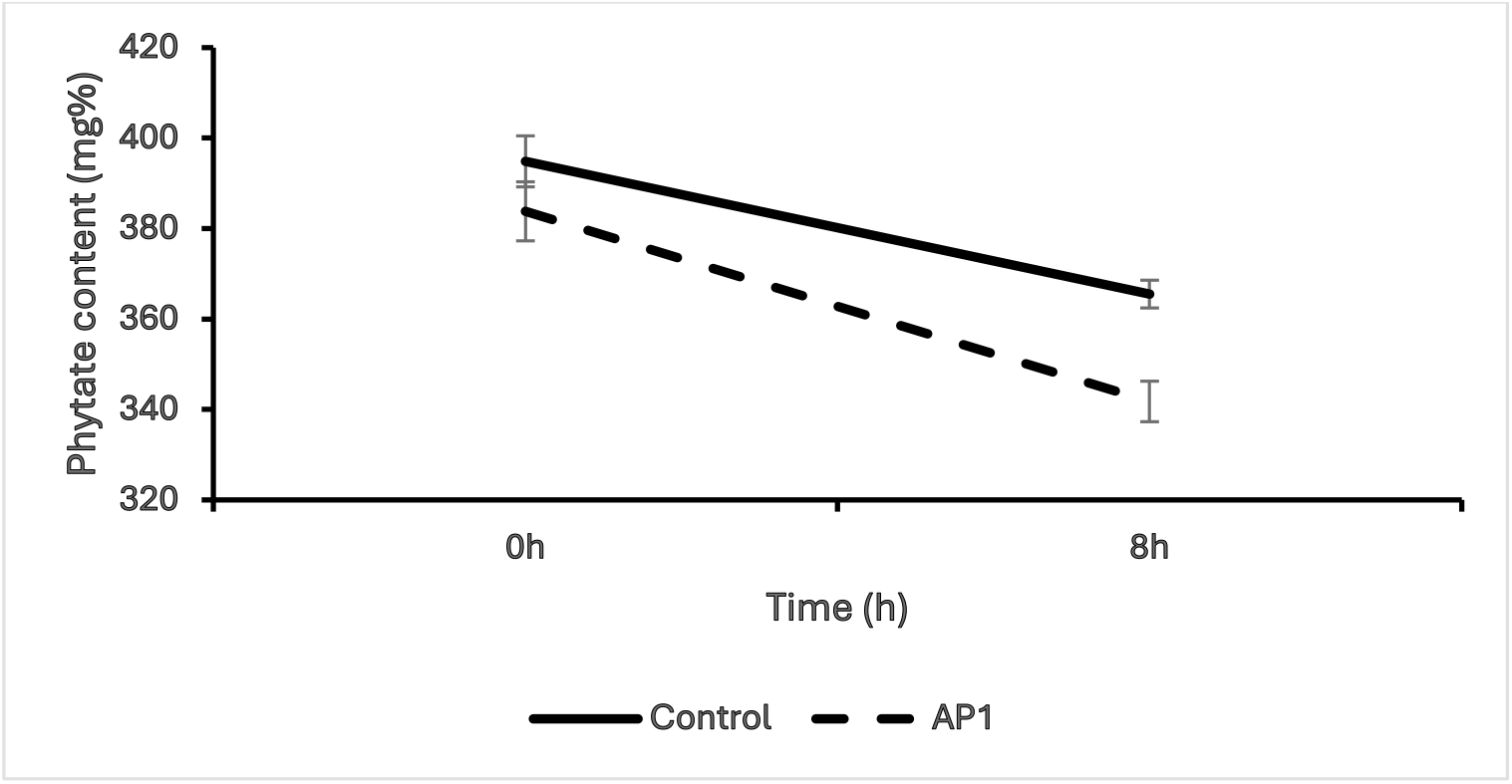
Reduction of phytate content in sorghum flour after 8 h of proofing with control and AP1 strains.

## 4 Discussion

A number of strategies have been devised to reduce phytate content in grains, including milling, soaking, fermentation and germination. These methods for phytate reduction have drawbacks, including loss of minerals and reduction of protein content (Gupta et al., 2015). In case of fermentation and germination, the extent of phytate reduction has been reported to be greater; however, the process is time consuming (approximately 2-3 days), which is a major drawback from an industrial perspective.

A widely used strategy for phytate reduction involves treatment with phytase enzyme (Gupta et al., 2015). Phytase derived from microbial sources is the most commonly used exogenous enzyme in the feed for monogastric animals and poultry (Dersjant-Li et al., 2015). A number of studies have been carried out to determine the effect of phytases on phosphate utilization and excretion, and a lot of effort has been expended on developing phytases that can survive the low pH of the stomach (Dersjant-Li et al., 2015).

Transgenic expression of phytases in crop plants (Abid et al., 2017) and downregulation of genes involved in phytic acid biosynthesis (Aggarwal et al., 2018) have been reported to increase bioavailability of iron and zinc (Abid et al., 2017; Aggarwal et al., 2018). However, transgenic crops may face regulatory hurdles and may have low acceptability.

Phytase has been expressed in the yeast *Saccharomyces cerevisiae* for improved bioethanol production using raw material used in distilleries (Mikulski et al., 2014; Chen et al., 2016; Mikulski et al., 2017). Phytase-producing yeast strains, including mutant *S. cerevisiae* and a natural *Pichia kudriavzevii* isolate, have been reported to reduce phytate content in wheat/cassava/sorghum flour (Vilanculos et al., 2023).

The presence of phytate in sorghum causes low mineral availability (Wu et al., 2018), limiting its attractiveness as an alternative to wheat for producing gluten-free foods. In the present study, the phytase gene from *A. japonicus* was integrated in the *S. cerevisiae* genome and expressed extracellularly. The constructed strain was used to reduce phytate content in sorghum dough, while simultaneously proofing it. This approach can be used along with other established procedures, such as adding exogenous phytase enzyme as well as germination, to further reduce phytate content in sorghum and other millets, thus enhancing their nutritional value.

## Acknowledgement

Rutuja Kamble was supported by a fellowship from Department of Biotechnology, Government of India, for the MTech Bioprocess Technology programme at Institute of Chemical Technology.

